# Chemokine induces phase transition from non-directional to directional migration during angiogenesis

**DOI:** 10.1101/2025.01.06.631616

**Authors:** Ning Gui, Keisuke Sako, Moe Fukumoto, Naoki Mochizuki, Hiroyuki Nakajima

## Abstract

During angiogenesis, sprouting endothelial cells (ECs) migrate and eventually connect to target vessels to form new vessel branches. However, it remains unclear how these sprouting vessels migrate toward the target vessels in three-dimensional space. We performed *in vivo* imaging of the cerebral capillary network formation in zebrafish to investigate how sprouting tip cells migrate toward their targets. Of note, we found that tip cells reach the target vessels through two phases: a non-directional phase and a directional phase. In the non-directional phase, sprouting tip cells dynamically extend and retract their protrusions at the leading front and have less directionality in their movement. In contrast, once tip cells enter the directional phase, they migrate directly toward the anastomotic targets. Chemokine receptor Cxcr4a and its ligand Cxcl12b are important for the phase transition to the directional phase. In *cxcr4a* mutants, sprouting tip cells lose their directionality and tend to connect to nearby sprouting ECs, resulting in altered capillary network patterning. Furthermore, in wild-type (WT) larvae, local Ca^2+^ oscillations were detected in protrusions of tip cells, specifically in the non-directional phase, but almost disappeared in the directional phase as a result of the Cxcr4-dependent phase transition. Thus, this study provides evidence of a chemokine-induced phase transition in migrating tip cells, which is important for proper vascular network formation in the zebrafish brain.

## Introduction

The vascular network is crucial in vertebrates for delivering oxygen and nutrients to tissues throughout the body. In development to adulthood, branched structures of blood vessels form mainly by sprouting angiogenesis, which involves the emergence of new vessels from the pre-existing ones (Eilken and Adams, 2010; Herbert and Stainier, 2011; Huveneers and Phng, 2024). Angiogenesis consists of multiple steps (Betz *et al*., 2016). First, an endothelial cell (EC) is selected from a pre-existing vessel in response to angiogenic cues such as vascular endothelial growth factor (VEGF)-A to become a migratory tip cell. As the tip cell migrates outward from the parental vessel and trailing stalk cells follow, the sprout elongates in contact with pre-existing vessels (Gerhardt *et al*., 2003). Endothelial lumen forms and expands in the sprout (Gebala *et al*., 2016). Eventually, the sprouts connect with the target vessels via anastomosis. Behaviors of ECs at each step of angiogenesis have been extensively studied (Eilken and Adams, 2010; Hogan and Schulte-Merker, 2017). However, little is known about how the sprout (tip cell) migrates toward the anastomotic targets in three-dimensional space.

Tip cells migrate by extending membrane protrusions containing actin-rich filopodia and lamellipodia at the leading edge. Among them, filopodia are thought to function as sensors of guidance cues (Huveneers and Phng, 2024; Zakirov *et al*., 2021). VEGF-A is required for filopodia extension (Gerhardt et al., 2003) and, thereby, has been considered to guide tip cell migration during sprouting angiogenesis. Besides VEGF-A, chemokines can also act as guidance cues. Especially, the chemokine receptor CXCR4 and its ligand CXCL12 (also known as SDF-1) play an important role in EC migration during angiogenesis (Kiefer and Siekmann, 2011). In mice, loss of CXCR4 and CXCL12 show defects in vascular development of various organs, such as the kidney (Takabatake *et al*., 2009), lung (Chandrasekaran *et al*., 2022), and retina (Pitulescu *et al*., 2017). In zebrafish, Cxcr4a (an ortholog of CXCR4) is expressed in sprouting ECs and has a role in guiding the formation of the coronary artery (Harrison *et al*., 2015; Ivins *et al*., 2015) and eye vasculature during development (Hasan *et al*., 2017), and vascular plexus formation during fin regeneration (Harrison et al., 2015; Ivins et al., 2015). At earlier stages of zebrafish cerebral vascular formation (around 36 hours post fertilization (hpf)), Cxcl12b, expressed along the midline of the neural keel, immediately above the basilar artery (BA), guides the sprout to connect to the BA (Bussmann *et al*., 2011). However, it remains uncertain how such guidance cues regulate the migratory behavior of sprouting ECs toward the anastomotic targets.

In the present study, we performed *in vivo* time-lapse imaging of cerebral capillary network formation in zebrafish. We show that sprouting tip cells undergo two distinct phases: a non-directional phase and a directional phase, in which they exhibit different cellular behaviors. Cxcr4a and Cxcl12b are important for inducing the phase transition to the directional phase, thereby promoting directional migration toward the anastomotic target. Finally, we show that Cxcr4a-mediated directional migration is important for the formation of proper capillary network formation.

## Materials and Methods

### Zebrafish husbandry and strains

Zebrafish of the AB strain (Danio rerio) were maintained and bred in 28 °C water (pH 7.25 and conductivity 500 μS) with a 14 h on/10 h off light cycle. Embryos and larvae were incubated at 28 °C in E3 medium. All zebrafish husbandry was performed under standard conditions according to institutional (National Cerebral and Cardiovascular Center) and national (Japan) ethical and animal welfare regulations. The experiments using zebrafish were approved by the animal committee of the National Cerebral and Cardiovascular Center (No.22054) and performed according to our institutional regulation.

The following transgenic and mutant zebrafish lines were used for this study: *Tg(kdrl:EGFP-CAAX,crya:EGFP)^ubs47^* (Heutschi *et al*., 2023), *Tg(kdrl:EGFP)^s843^* (Jin *et al*., 2005), *Tg(fli1:H2B-mCherry)^ncv31^* (Yokota *et al*., 2015), *Tg(UAS:GCaMP7a)* (Muto *et al*., 2013), *Tg(kdrl:TagBFP)^mu293^* (Matsuoka *et al*., 2016), *Cxcr4a^s421^* (Schmid *et al*., 2013) and *Tg(fli1:Gal4FF)* (Herwig *et al*., 2011). Throughout the text, all Tg lines used in this study are simply described without their line numbers. For example, *Tg(fli1:H2B-mCherry)^ncv31^* is abbreviated to *Tg(fli1:H2B-mCherry)*.

The full locus deletion alleles ncv148 for *cxcl12a* and ncv149 for *cxcl12b* genes were generated by CRISPR-Cas9 techniques as described below.

### Generation of full locus deletion mutants by CRISPR-Cas9 system

To generate full locus deletion alleles of *cxcl12a* and *cxcl12b*, double guide RNAs (gRNAs) were designed upstream of the 5’ UTR and downstream of the 3’ UTR, flanking the region to be deleted (see Figure 3A). crRNA sequences were designed using the CRISPR direct software (https://crispr.dbcls.jp) as follows: *cxcl12a*-5’-guide: 5’-AGTGTGAACTCTCCCACCGA-3’, *cxcl12a*-3’-guide: 5’-GAAGACTTAAATTCACACCT-3’, *cxcl12b*-5’-guide: 5’-ATGTTGAAGAATGCTATCGT-3’, *cxcl12b*-3’-guide: 5’-TAGGCCAGGTTCTGTCCCGG-3’. crRNAs and tracrRNA were generated by IDT Inc, and the RNP complex was prepared according to the manufacturer’s instructions.

Embryos, injected with the RNP complex at the one-cell stage, were raised to adulthood and crossed with AB to identify full locus deletion founders. Screening for founders was conducted by genomic PCR and subsequent sequencing using the primer sets shown in Table S1. For the genotyping of the mutants, PCR analyses of genomic DNAs were routinely performed using the following three primer sets: cxcl12a(b)-5’-Fw and cxcl12a(b)-5’-Rev for WT allele, cxcl12a(b)-3’-Fw and cxcl12a(b)-3’-Rev for WT allele, cxcl12a(b)-5’-Fw and cxcl12a(b)-3’-Rev for full locus deletion allele. This allows us to distinguish between WT, Het, and Homo.

### *cxcr4a* crispants

The double-target CRISPR-mediated F0 knockout (Chiba *et al*., 2024) was generated by the sgRNA (IDT Inc.) targeting the following sequence: 5′-GGACATCGGAGCCAACTTTG -3′, 5′-CTGCGAGCGCATATACCCGC-3′. The preparation of RNP complex was prepared according to the manufacturer’s instructions, except that 0.7 μl of each crRNA was used to prepare the gRNA solution.

### Chemical treatment

*Tg(kdrl:EGFP)* larvae were treated with 1 μM ki8751, an inhibitor of Vegfr2, at 2 h after the start of confocal imaging.

### Image acquisition and processing

The pigmentation of embryos and larvae was inhibited by treatment with 1-phenyl-2-thiourea (PTU) (Sigma-Aldrich). For confocal imaging, zebrafish embryos were dechorionated, anesthetized in 0.016% tricaine (Sigma-Aldrich) in E3 medium, and mounted in 1% low-melting agarose poured onto a 35-mm diameter glass-based dish (Asahi Techno Glass). Confocal imaging in Fig. 5D, S1A, S3A, S4B and Movie 3 and 4 was performed on Andor Dragonfly spinning disc confocal (Andor Technology Ltd) based on ECLIPSE FN1 upright microscope (Nikon), equipped with water-immersion LWD 16x/0.80 NA (Nikon), Zyla4.2 PLUS USB 3.0 sCMOS cameras (Andor Technology Ltd) and P-725.4 PIFOC piezo nano-positioner (Physik Instrumente) regulated with Fusion software (Andor Technology Ltd). Other confocal images were taken with a FluoView FV1200 or FV3000 confocal upright microscope system (Olympus) equipped with water-immersion XLUMPlan FL N 20x/1.00 NA (Olympus) and a multi-alkali or GaAsP photomultiplier tube regulated with FluoViewASW software (Olympus). The 405 nm, 473 nm, and 559 nm laser lines were used. Images were acquired sequentially to avoid cross-detection of the fluorescent signals.

All confocal images were processed and analyzed with IMARIS 9.5.1, 9.9.1, or 10.1.0 software (Oxford Instruments). In Fig. 1C, 4A, and 4B, and Movie 1, *kdrl:TagBFP^+^* EC regions were extracted by the masking function to exclude non-specific signals. In Fig. 1D, the outline of the nucleus at each time point was automatically extracted.

**Figure 1.**
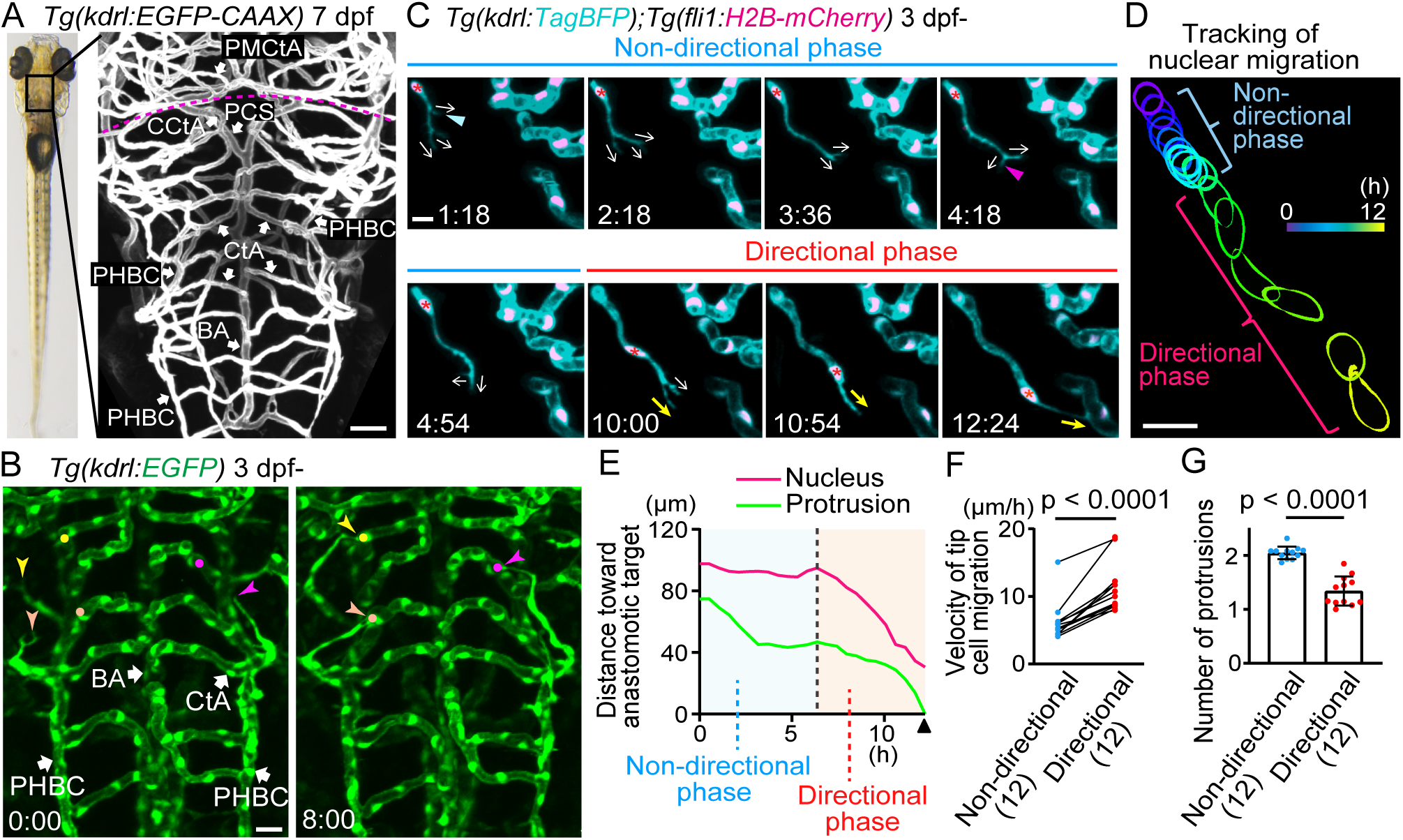
Sprouting tip cells undergo two phases, non-directional and directional phases, to reach the target vessels. (A) Representative confocal image of the brain of a *Tg(kdrl:EGFP-CAAX)* larva (7 dpf). Plasma membrane of endothelial cells (ECs) is labeled with EGFP-CAAX. Dorsal view, anterior to the top. The dotted line indicates the boundary between the hindbrain (posterior) and midbrain (anterior). (B) Time-sequential images of a *Tg(kdrl:EGFP)* larva (from 3 dpf). Elapsed time (h:min). *kdrl*:EGFP^+^ ECs are shown as green. ECs sprouting (arrowheads) from the primordial hindbrain channel (PHBC) eventually connect to the central artery (CtA) (circles). (C) Time-sequential images of a *Tg(kdrl:TagBFP);Tg(fli1:H2B-mCherry)* larva (from 3 dpf). *kdrl*:TagBFP*^+^* ECs and *fli1*:*H2B-*mCherry^+^ EC nuclei are shown as cyan and magenta, respectively. Direction of protrusions is indicated by arrows. In the non-directional phase, tip cells sprouting from the PHBC dynamically retract (blue arrowhead) and extend (magenta arrowhead) their protrusions in various directions (arrows). Then, the nuclei of tip cells (asterisks) rapidly move toward the anastomotic targets, the CtAs, in the directional phase. (D) Tracking of the nucleus of the tip cell shown in (C). The outline of the nucleus is depicted every 0.5 h as different colors. (E) Quantitative analysis of the data shown in (C). Minimum distance between the anastomotic target and the tip cell nucleus (magenta) or the distal edge of the protrusion (green) at each time point until the connection to the target vessel (arrowhead). (F) Graph shows the time-averaged velocity of the tip cell nucleus in the non-directional and directional phases. Data are mean ± s.d. (n = 12 cells from 6 larvae). (G) Graph shows the time-averaged number of protrusions per tip cell throughout the non-directional and directional phases. Data are mean ± s.d. (n = 12 cells from 6 larvae). Scale bar:10 μm. *P* values were determined by paired (F) or unpaired (G) two-tailed Student’s t-test. BA, basilar artery; PHBC, primordial hindbrain channel; CtA, central artery; CCtA, cerebellar central artery; PCS, posterior communicating segment; PMCtA, posterior metencephalic central artery.

### *in vivo* Ca^2+^ imaging of sprouting tip cells

For long-term imaging for GCaMP7a signals in Fig. 4A and 4B, we performed 3D time-lapse imaging using an FV3000 confocal microscope every 5 min. Under these experimental conditions, we could capture GCaMP7a signals only when the region of interest was scanned and could not capture all local Ca^2+^ oscillations because the duration of each oscillation was around 20 sec. Therefore, to visualize local Ca^2+^ oscillations in Fig. S3A and S4B, we performed high-speed 3D time-lapse imaging using a Dragonfly spinning disc confocal microscope every 15 sec with a z-interval of 6 μm. Especially, to quantify the frequency of local Ca^2+^ oscillations in Fig. 4C, 3D images were taken every 15 sec for 10 min. Z-stack images were 3D volume-rendered and analyzed with IMARIS 9.5.1, 9.9.1, or 10.1.0 software (Oxford Instruments).

### Quantitative analyses

For quantification of Fig. 1E, spots were manually created at the center of tip cell nucleus, the distal edge of the protrusion, and the target site of anastomotic vessel at each time point. The minimum distance from the spot of the target site to the spot of the nucleus or protrusion was measured at each time point.

For quantification of Fig. 2D, a spot was manually created at the center of tip cell nucleus at each time point. The minimum distance from the spot of the nucleus at the indicated time point to the spot of the nucleus at the end time point (at 12 h) was measured at each time point.

**Figure 2.**
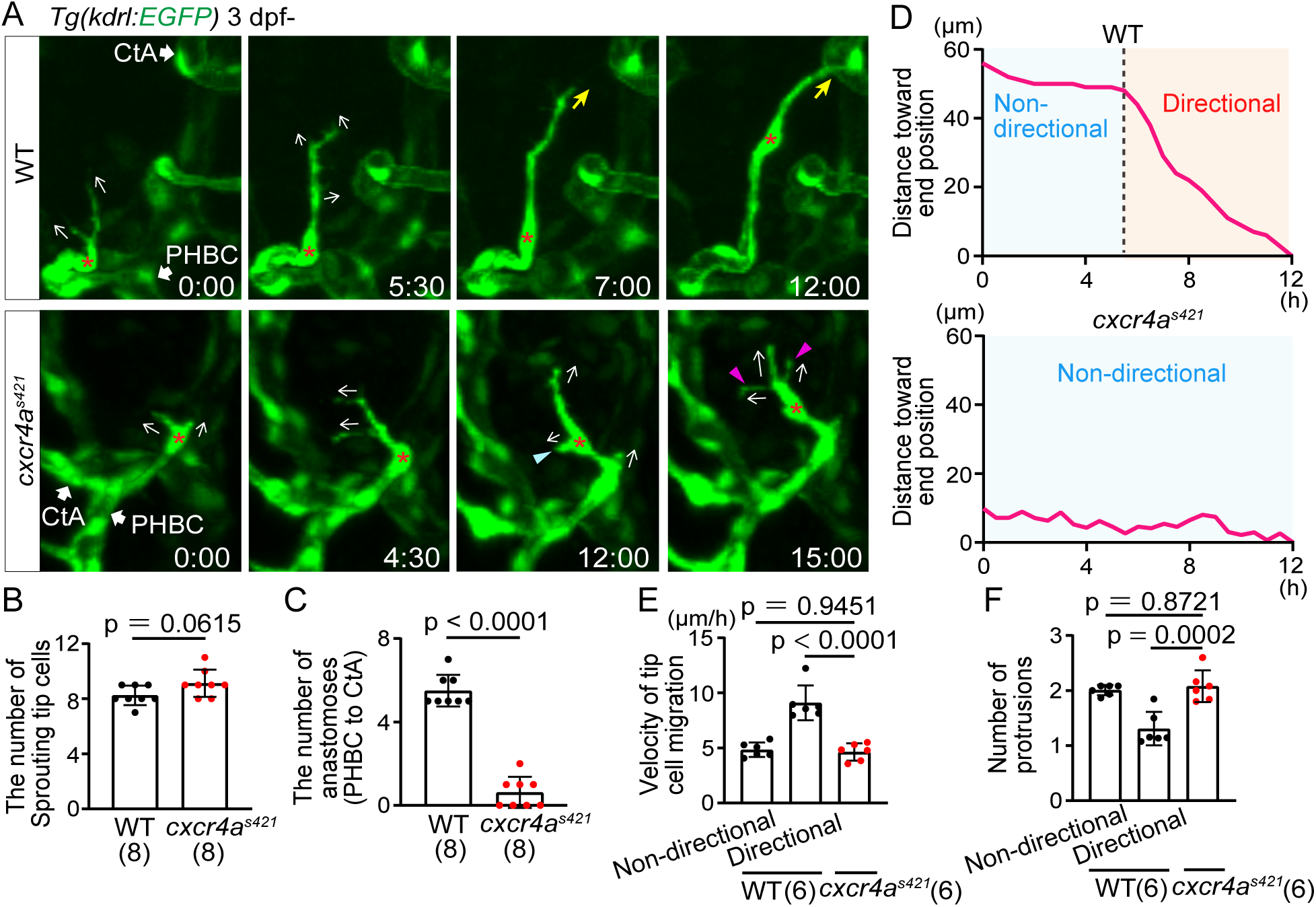
Cxcr4a is important for directional migration to anastomotic targets. (A) Time-sequential images of *Tg(kdrl:EGFP)* WT and *cxcr4a^s421^* sibling larvae (from 3 dpf). Asterisks indicate tip cell nuclei. In *cxcr4a^s421^* larvae, tip cells sprouting from the PHBC repeatedly extend (magenta arrowheads) and retract (blue arrowhead) their protrusions in various directions (arrows). They do not migrate toward the target vessel. (B) Graph shows the number of tip cells sprouting from the PHBC in WT and *cxcr4a^s421^* sibling larvae (from 73 hpf to 121 hpf). Data are mean ± s.d. (WT, n = 8 larvae; *cxcr4a^s421^*, n = 8 larvae). Each dot represents an individual larva in this graph. (C) Graph shows the number of connections of tip cells sprouting from the PHBC to the CtAs or CCtAs in WT and *cxcr4a^s421^* sibling larvae (from 73 hpf to 121 hpf). Data are mean ± s.d. (WT, n = 8 larvae; *cxcr4a^s421^*, n = 8 larvae). (D) Quantitative analysis of the data shown in (A). Minimum distance between the position of the tip cell nucleus at each time point and the position of the nucleus at 12 h in WT and *cxcr4a^s421^* sibling larvae. Connection to the target CtA occurs around 12 h in this WT larva as shown in (A). (E) Graph shows the time-averaged velocity of the tip cell nucleus in WT (non-directional and directional phases) and *cxcr4a^s421^* sibling larvae. Data are mean ± s.d. (WT, n = 6 cells in 4 larvae; *cxcr4a^s421^*, n = 6 cells in 4 larvae). (F) Graph shows the time-averaged number of protrusions per tip cell in WT (non-directional and directional phases) and *cxcr4a^s421^* sibling larvae. Data are mean ± s.d. (WT, n = 6 cells in 4 larvae; *cxcr4a^s421^*, n = 6 cells in 4 larvae). Scale bar:10 μm. *P* values were determined by two-tailed Student’s t-test (B, C) and one-way ANOVA with Tukey’s test (E,F).

For quantification of Fig. 1F and 2E, the velocity of tip cell was measured by tracking the nucleus as in Fig. 1E. The time-averaged velocity of each larva is shown as each dot. For quantification of Fig. 1G and 2F, the time-averaged protrusion number was measured and shown as each dot. In 13.3% of cases, as seen in Fig. S1B, directional migration stopped until the onset of stalk cell migration. Such cases were excluded from the quantitative analyses between the non-directional and directional phases as they were not typical directional phases.

For quantification of Fig. 2B, 2C, 3E and 3F, the number of sprouting tip cells from the PHBC and the number of anastomoses from the PHBC to CtA were quantified from 73 hpf (3 dpf) to 121 hpf (5 dpf) for 48 h. Quantification was performed in the whole hindbrain.

**Figure 3.**
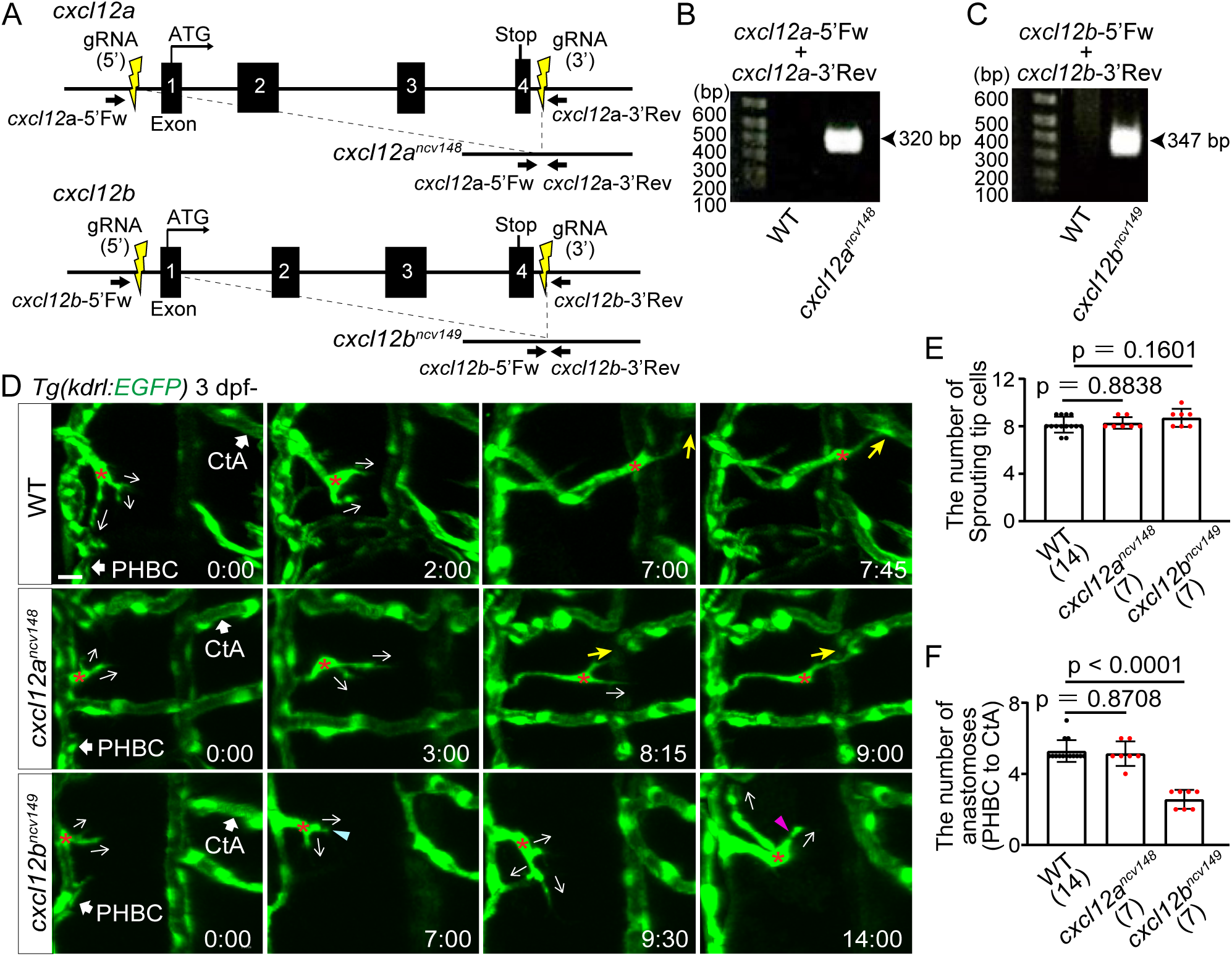
Cxcl12b but not Cxcl12a is responsible for directional migration to anastomotic targets. (A) Schematic representation for generating full locus deletion alleles in *cxcl12a* and *cxcl12b*. Double guide RNAs (gRNAs) (yellow) were designed upstream of the 5’ UTR and downstream of the 3’ UTR, flanking all exons. (B,C) Images of gels showing full locus deletions in *cxcr12a^ncv148^* (B) and *cxcr12b^ncv149^* homozygous mutants (C). Primer sites are shown in (A). (D) Time-sequential images of *Tg(kdrl:EGFP)* WT, *cxcr12a^ncv148^*, and *cxcr12b^ncv149^* larvae (from 3 dpf). Asterisks indicate tip cell nuclei. In WT and *cxcr12a^ncv148^* larvae, tip cells sprouting from the PHBC finally undergo directional migration to the target CtAs (yellow arrows). In contrast, in *cxcr12b^ncv149^* larvae, tip cells sprouting from the PHBC repeatedly extend (magenta arrowhead) and retract (blue arrowhead) their protrusions while extending them in various directions (arrows), and do not migrate toward the target. (E) Graph shows the number of tip cells sprouting from the PHBC in WT, *cxcr12a^ncv148^*, and *cxcr12b^ncv149^* larvae (from 73 hpf to 121 hpf). Data are mean ± s.d. (WT, n = 14 larvae; *cxcr12a^ncv148^*, n = 7 larvae; *cxcr12b^ncv149^* , n = 7 larvae). (F) Graph shows the number of connections of tip cells sprouting from the PHBC to the CtAs or CCtAs in WT, *cxcr12a^ncv148^*, and *cxcr12b^ncv149^* larvae (from 73 hpf to 121 hpf). Data are mean ± s.d. (WT, n = 14 larvae; *cxcr12a^ncv148^*, n = 7 larvae; *cxcr12b^ncv149^* , n = 7 larvae). Scale bar:10 μm. *P* values were determined by one-way ANOVA with Tukey’s test.

For quantification of Fig. 5B and 5C, the number of anastomoses was quantified from 73 hpf (3 dpf) to 121 hpf (5 dpf) in the whole hindbrain. For quantification of Fig. 5E, the number of small loops less than 20 μm was counted in the CtA network of the whole hindbrain by setting a sphere with a radius of 20 μm using IMARIS software.

### Data analysis and statistics

Data were analyzed using GraphPad Prism software or Excel and were presented as mean ± s. d. Sample numbers and the statistical methods are indicated in figure legends.

## Results

### Sprouting tip cells undergo two phases, non-directional and directional phases, to reach the anastomotic targets

To visualize how sprouting vessels migrate to their target vessels, we here focused on the formation of the hindbrain capillary network in zebrafish. In the hindbrain, the central arteries (CtAs) and cerebellar central arteries (CCtAs) form a capillary network (Fig. 1A) (Isogai *et al*., 2001), which creates a CNS-specific blood-brain barrier (Xie *et al*., 2010). Both the CtAs and CCtAs are derived from sprouting from the primordial hindbrain channels (PHBC) located bilaterally, and connect to the BA and the posterior communicating segment (PCS) respectively at earlier stages of the cerebral vascular development (∼1.5 days post fertilization (dpf)) (Bussmann et al., 2011; Chen *et al*., 2019; Isogai et al., 2001). For simplicity, the CtA and CCtA are hereafter referred to as the CtA. At later stages (after 3 dpf), sprouts from the PHBC connect to existing CtAs, thereby forming a capillary network of CtAs (Fig. 1B). Finally, CtA networks show different patterns in different individuals (Fig. S1A).

As a model of capillary network formation, we carefully observed how sprouts from the PHBC connect to the CtAs by time-lapse imaging. Our imaging analyses have revealed that sprouting ECs undergo two distinct phases before connecting to the target vessels (Fig. 1C-1G). Initially, sprouting tip cells extended and retracted multiple protrusions (average number = 2.05, Fig. 1G) in various directions (Fig. 1C) and had less directionality toward the anastomotic targets (Fig. 1C-1E, Movie 1). We here define this phase as a “non-directional phase.” After this phase, the tip cells rapidly migrated toward the target vessels, the CtAs (Fig. 1C-1F), while their protrusions were mostly directed to their targets (Fig. 1C: yellow arrows, Movie 1). We define this phase as a “directional phase.” Our quantification analyses revealed that the average migration velocity of tip cell nuclei was significantly faster in the directional phase than in the non-directional phase (Fig. 1F). In addition, the time-averaged number of protrusions became significantly less in the directional phase (average number = 1.34, Fig. 1G). Therefore, our results demonstrate that during the formation of the zebrafish cerebral capillary network, sprouting tip cells undergo two distinct phases that exhibit different cellular behavior.

We also found that, in 13.3% of the cases, directional migration of tip cells stopped before arrival to the anastomotic targets. In such cases, directional migration restarted after stalk cells migrate out of the parental vessel (Fig. S1B). These results suggest that coordinated migration of tip and stalk cells is also important for the directional migration of angiogenic sprouts.

### Cxcr4a is required for directional migration to anastomotic targets

Next, we examined which extracellular stimuli regulate the transition from the non-directional to directional phase. VEGF-A is a known guidance cue for sprouting ECs (Gerhardt et al., 2003). When we suppressed VEGF-A/VEGFR2 signaling by the treatment with ki8751, an inhibitor of Vegfr2 (also known as Kdrl in zebrafish) (Yokota et al., 2015), the initial tip cell sprouting from the PHBC was completely blocked, and budding sprouts were retracted to the PHBC (Fig. S2). These results suggest that VEGF is important for the initial tip cell sprouting from the PHBC. However, considering that VEGF-A/VEGFR2 signaling regulates multiple aspects of angiogenesis, including tip cell selection, sprouting, migration, and proliferation (Koch and Claesson-Welsh, 2012; Lohela *et al*., 2009), it is technically difficult to segregate the function of VEGF-A as a guidance cue.

CXCL12/CXCR4 signaling is another guidance cue for ECs during angiogenesis (Kiefer and Siekmann, 2011). To examine the role of the CXCL12-CXCR4 axis, we performed time-lapse imaging of *cxcr4a* mutants, focusing on the connection of tip cells from the PHBC to the CtA. In *cxcr4a* mutants, tip cells sprouted from the PHBC and extended protrusions similarly to WT (Fig. 2A). The number of tip cells sprouting from the PHBC in *cxcr4a* mutants was also comparable to WT (Fig. 2B). These results suggest that tip cells acquire angiogenic capacity even in *cxcr4a* mutants, presumably by VEGF-A/VEGFR2 signaling. However, most of these sprouting ECs could not connect to the CtAs in *cxcr4a* mutants (Fig. 2A and 2C). In these mutants, tip cells repeatedly extended and retracted their protrusions in various directions (Fig. 2A), but lost directionality in their movement (Fig. 2A and 2D, Movie 2). In the tip cells of *cxcr4a* mutant larvae, the velocity and the number of protrusions are comparable to those in the non-directional phase of WT (Fig. 2E and 2F), suggesting that tip cells in the *cxcr4a* mutants persistently retain the cellular characteristics of the non-directional phase.

Therefore, our results indicate that Cxcr4a is important for the transition from the non-directional phase to the directional phase.

### Cxcl12b but not Cxcl12a is responsible for directional migration to anastomotic targets

Next, we investigated the guidance cues for the directional EC migration. As orthologues of CXCL12, zebrafish have Cxcl12a and Cxcl12b, both of which act as ligands for Cxcr4a in a context-dependent manner (Bussmann et al., 2011; Xu *et al*., 2014). To examine the roles of Cxcl12a and Cxcl12b, we analyzed the effects of loss of these ligands. Here, to avoid genetic compensation (El-Brolosy *et al*., 2019), we established full locus deletion fish of *cxcl12a* and *cxcl12b* (Fig. 3A). All exons are deleted in these mutant alleles (Fig. 3B and 3C). In both *cxcl12a* mutants and *cxcl12b* mutants, tip cells sprouted normally from the PHBC, in numbers comparable to WT (Fig.3D and 3E). These sprouting tip cells were normally connected to the CtAs in *cxcl12a* mutants as in WT (Fig. 3D and 3F). In contrast, in *cxcl12b* mutants, about half of tip cells did not connect to the CtA, and persistently extended and retracted their protrusions (Fig. 3D and 3F). Thus, *cxcl12b* mutant larvae but not *cxcl12a* mutants phenocopied *cxcr4a* mutants, although the effects were slightly milder. These results strongly suggest that Cxcl12b acts as a guidance cue to control directional migration toward anastomotic targets during cerebral capillary network formation.

### Local Ca^2+^ oscillations occur in protrusions at the leading front in the non-directional phase

We previously showed that intracellular Ca^2+^ dynamics reflect angiogenic capacity in tip cells sprouting from the dorsal aorta (DA) (Yokota et al., 2015). To investigate the Ca^2+^ dynamics during non-directional and directional migration of tip cells, we conducted *in vivo* Ca^2+^ imaging by expressing GCaMP7a (GFP-based Ca^2+^ probe) in ECs. Of note, in addition to Ca^2+^ oscillations occurring throughout the cell (see Fig. S3A) (Yokota et al., 2015), local Ca^2+^ oscillations were detected in protrusions at the leading front of sprouting tip cells in the non-directional phase (Fig. 4A and S3A). In contrast, these local Ca^2+^ oscillations mostly disappeared after the transition to the directional phase (Fig. 4A and S3A). We quantified these local Ca^2+^ oscillations and found that oscillation frequency was markedly reduced in the directional phase (Fig. 4C).

**Figure 4.**
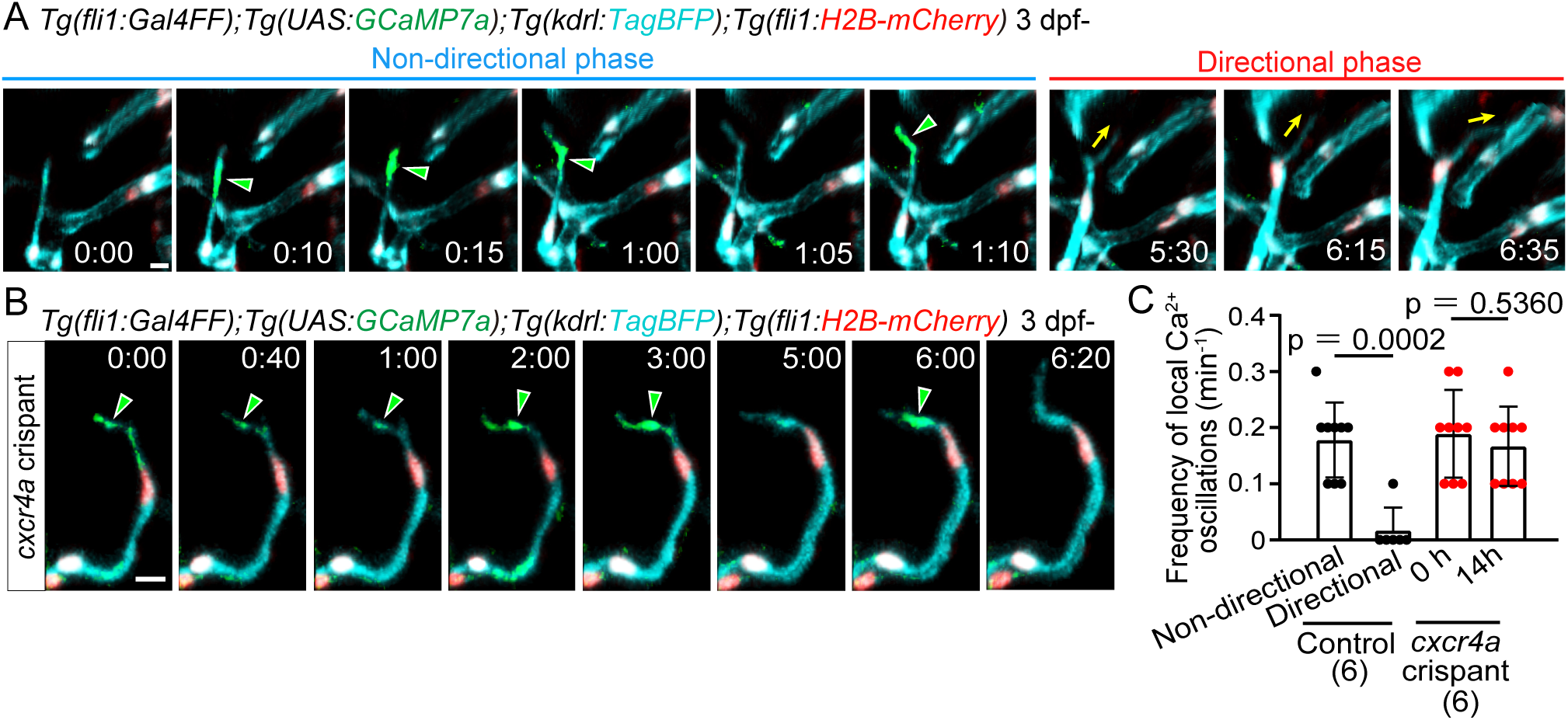
Local Ca^2+^ oscillations occur specifically in the non-directional phase. (A) Time-sequential images of a *Tg(fli1:Gal4FF);Tg(UAS:GCaMP7a);Tg(kdrl:TagBFP);Tg(fli1:H2B-mC)* larva (3 dpf) after tip cell sprouting from the PHBC. Elapsed time (h:min). Local Ca^2+^ signals (green arrowheads) are detected in the leading front of tip cells in the non-directional phase, but rarely in the directional phase. (B) Time-sequential images of a *Tg(fli1:Gal4FF);Tg(UAS:GCaMP7a);Tg(kdrl:TagBFP);Tg(fli1:H2B-mC) cxcr4a* crispant larva (F0 injected larva) (3 dpf) after tip cell sprouting from the PHBC. Local Ca^2+^ signals are maintained in the leading front of tip cells (green arrowheads) that do not head to the target vessels. (C) Graph shows the frequency of local Ca^2+^ oscillations in the leading front of tip cells in control and *cxcr4a* crispants (3-4 dpf). Data are mean ± s.d. (WT: n = 9 cells in the non-directional phase, n = 6 cells in the directional phase in 6 larvae; *cxcr4a^s421^*: n = 9 cells (at 0 h), 9 cells (at 14 h) in 6 larvae). Scale bar:10 μm. *P* value was determined by unpaired two-tailed Student’s t-test.

Next, to investigate whether these changes in calcium dynamics are related to the Cxcr4a-dependent phase transition, we examined the effect of loss of Cxcr4a on the local calcium dynamics. We observed *cxcr4a* F0 knockout zebrafish injected with *cxcr4a* gRNAs (called *cxcr4a* crispants). As in the *cxcr4a* mutants, in *cxcr4a* crispants, sprouting tip cells extended and retracted their protrusions, and have less directionality toward the anastomotic targets (Fig. S4A). These larvae persistently exhibited local Ca^2+^ oscillations in their extending and retracting protrusions (Fig. 4B, and S4B). These results indicate that the Cxcr4a-dependent phase transition causes the disappearance of local Ca^2+^ oscillations in tip cells undergoing directional migration.

Previously, similar local Ca^2+^ oscillations were reported in retracting protrusions of tip cells (Liu *et al*., 2020). In our analysis, local Ca^2+^ oscillations were observed not only in retracting protrusions but also in extending protrusions (Fig. S3B). These results suggest that local Ca^2+^ oscillations might be involved in the dynamic extension and retraction of unstable tip cell protrusions.

### Cerebral capillary network patterning becomes altered in *cxcr4a* mutants

Finally, to delineate the role of Cxcr4a-mediated directional migration, the phenotype of *cxcr4a* mutants in cerebral capillary network formation was carefully examined. In WT, sprouts from the PHBC anastomosed with the CtAs (Fig. 1B and 1C), whereas in *cxcr4a* mutants, most sprouts failed to connect with the CtAs and remained as sprouts (Fig. 2A and 2C). We observed that some of these sprouts connected with the other sprouts from the nearby PHBC, although this did not occur in WT (Fig. 5A and 5B). In the cxcr4a mutants, there was increased sprouting from CtAs (Fig. 5A and 5C). These sprouts anastomosed with nearby sprouting ECs rather than with lumenized vessels (Figs 5A and 5C). Thus, in the *cxcr4a* mutants, anastomoses from sprouts to vessels, as seen in the WT (see Fig. 1B and 1C), were markedly reduced, whereas anastomoses from sprouts to sprouts were enhanced. As a result, more small vessel loops were formed in the cerebral capillaries in *cxcr4a* mutant (Fig. 5D and 5E). Thus, cerebral capillary network pattering becomes altered in *cxcr4a* mutants. These results suggest that Cxcr4a-mediated directional migration is important for the formation of proper capillary network formation.

**Figure 5.**
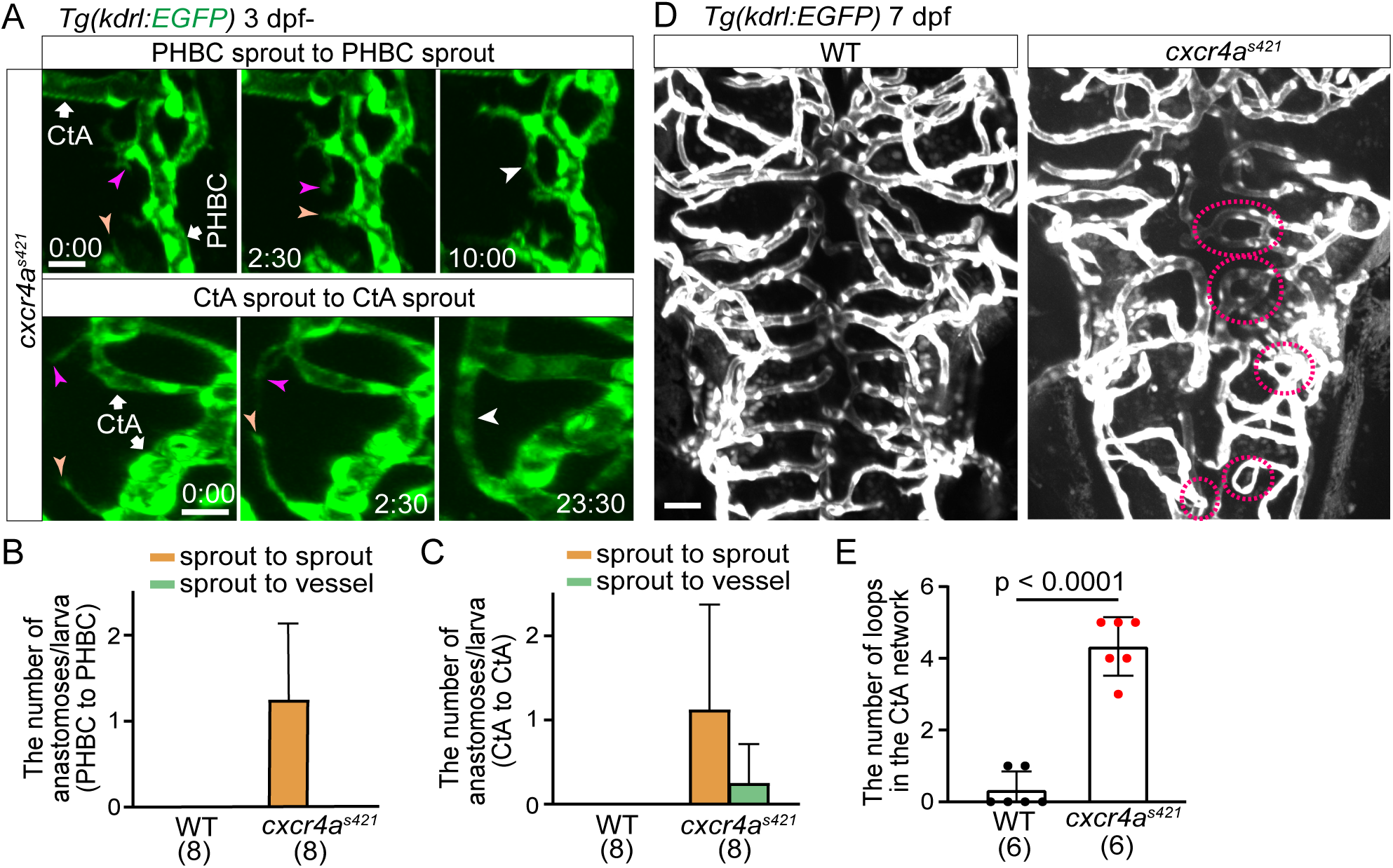
Cerebral capillary network patterning becomes altered in *cxcr4a^s421^* mutants. (A) Time-sequential images of *Tg(kdrl:EGFP) cxcr4a^s421^* mutant larva (from 3 dpf). Upper: a tip cell sprout from the PHBC (magenta arrowheads) connects with another tip cell sprout from the PHBC nearby (orange arrowheads). Lower: a tip cell ectopically sprouts from the CtA (magenta arrowheads) and is often connected with another tip cell sprouting from the nearby CtA (orange arrowheads). (B) Graph shows the number of tip cell sprouts from the PHBC anastomosing with other sprouts from the PHBC (sprout to sprout, orange bar) and the number of tip cell sprouts from the PHBC anastomosing with the PHBC vessel (sprout to vessel, green bar) in WT and *cxcr4a^s421^* sibling larvae (from 73 hpf to 121 hpf). Data are mean ± s.d. (WT, n = 8 larvae; *cxcr4a^s421^*, n = 8 larvae). (C) Graph shows the number of CtA sprouts anastomosing with other CtA sprouts (sprout to sprout, orange bar) or with CtA vessels (sprout to vessel, green bar) in WT and *cxcr4a^s421^* sibling larvae (from 73 hpf to 121 hpf). Data are mean ± s.d. (WT, n = 8 larvae; *cxcr4a^s421^*, n = 8 larvae). (D) *Tg(kdrl:EGFP)* WT and *cxcr4a^s421^* sibling larvae (7 dpf). Small vessel loops (dotted circles) are increased in *cxcr4a^s421^* mutants. (E) Quantitative analysis of the data shown in (D). The number of loops (less than 20 μm) in the CtA capillary network in the hindbrain of WT and *cxcr4a^s421^* sibling larvae. Data are mean ± s.d. (WT, n = 6 larvae; *cxcr4a^s421^*, n = 6 larvae). Scale bar:10 μm. *P* value was determined by unpaired two-tailed Student’s t-test.

## Discussion

In this paper, we found that tip cells reach the anastomotic targets through two distinct phases: the non-directional and directional phases. Whereas tip cells have less directionality in their movement in the non-directional phase, they migrate directly toward the anastomotic targets in the directional phase. Chemokine signaling Cxcl12b and Cxcr4a induce a phase transition from the non-directional phase to the directional phase. In *cxcr4a* mutants, sprouting tip cells lose their directionality and tend to connect to nearby sprouting ECs in the cerebral capillary formation. In contrast, Cxcr4a is dispensable for the formation of the intersegmental vessels (ISVs) in the zebrafish trunk (Hasan et al., 2017). In the ISV formation, sprouting ECs always run through a defined space between each pair of somites and eventually anastomose with neighboring sprouts on the dorsal side (Childs *et al*., 2002; Isogai *et al*., 2003). Therefore, Cxcr4a may not be required for angiogenesis through a pre-determined pathway. In this case, the location of Vegfa, a key driver of angiogenesis, and soluble Flt1 (sFlt1) and Semaphorin, which negatively act on VEGF-A signaling, determines the defined vascular patterns of the ISVs (Krueger et al., 2011; Yokota et al., 2015; Zygmunt et al., 2011). In contrast, in many other vascular beds, such as skin (Kam *et al*., 2023), heart (Harrison et al., 2015), retina (Gerhardt et al., 2003), and brain (Blinder *et al*., 2010), broad vascular patterns such as large arteries and veins are preserved, but the detailed branching patterns of capillaries differ among individuals. Because CXCR4/Cxcr4a mutant ECs generally show defective migration in such organs (Cavallero *et al*., 2015; Ivins et al., 2015; Li *et al*., 2013; Pitulescu et al., 2017), it is reasonable to speculate that zebrafish Cxcr4a and mammalian CXCR4 may commonly regulate directional migration of tip cells to form a capillary network without a defined pattern.

Our results here strongly suggest that Cxcl12b is a ligand for Cxcr4a to induce directional migration to anastomotic target. At earlier stages of zebrafish cerebral vascular formation (around 1.5 pdf), ECs sprout from the PHBC and connect to the centrally located BA to form the arch-shaped CtAs (Bussmann et al., 2011). At this stage, *cxcl12b* is expressed along the midline of the neural keel, immediately above the BA, and plays an essential role in targeting EC sprouts to reach the BA (Bussmann et al., 2011). On the other hand, *cxcl12b*-expressing cells have not been identified during the formation of the cerebral capillary network (3-5 dpf) when the PHBC-CtA connection occurs. If *cxcl12b*-expressing cells can be visualized, it is expected to reveal how anastomotic targets are determined to establish the proper cerebral capillary network in zebrafish.

*cxcr4a* mutants have been shown to have abnormal EC migration in various vascular beds (Bussmann et al., 2011; Harrison et al., 2015; Hasan et al., 2017; Xu et al., 2014). Our detailed time-lapse analyses of *cxcr4a* mutants revealed that Cxcr4a is dispensable for tip cell sprouting, including filopodia formation, but is required for directional migration toward the anastomotic target. On the other hand, our results and previous reports indicate that the initial tip cell sprouting is controlled by Vegfa/Vegfr2 signaling (Figure S2) (Ferrara, 2009; Lohela et al., 2009; Yokota et al., 2015). In summary, in sprouting angiogenesis in cerebral capillaries, Vegfa/Vegfr2 signaling first induces tip cell sprouting from the parental vessels. After sprouting, tip cells are angiogenic but have less directionality in the non-directional phase. Once tip cells enter the directional phase by Cxcr4a/Cxcl12b signaling, they migrate directly toward the anastomotic targets and finally produce the new vessel connections. Thus, we propose that Vegfa/Vegfr2-dependent tip cell sprouting and Cxcl12b/Cxcr4a-dependent tip cell targeting are distinct processes in sprouting angiogenesis.

## Abbreviations

EC: endothelial cell
VEGF: vascular endothelial growth factor
WT: wild-type
sgRNA: single guide RNA
crRNA: CRISPR RNA
RNP: ribonucleoprotein.

## Acknowledgments

We thank Didier Stainier (Max Planck Institute for Heart and Lung Research, Germany) for *cxcr4a^s421^*, Etienne Schmelzer and Heinz-Georg Belting (University of Basel, Switzerland) for *Tg(kdrl:EGFP-CAAX,crya:EGFP)^ubs47^*, and Akira Muto and Koichi Kawakami (National Institute of Genetics) for *Tg(UAS:GCaMP7a)* zebrafish. We are grateful to Arndt Siekmann (University of Pennsylvania) for helpful advice. We are grateful to M. Sone, T. Satoh, S. Toyoshima, and E. Hanimura for technical assistance.

