## Supplementary Materials for "Chemokine induces phase transition from non-directional to directional migration during angiogenesis"

### Supplementary Figures

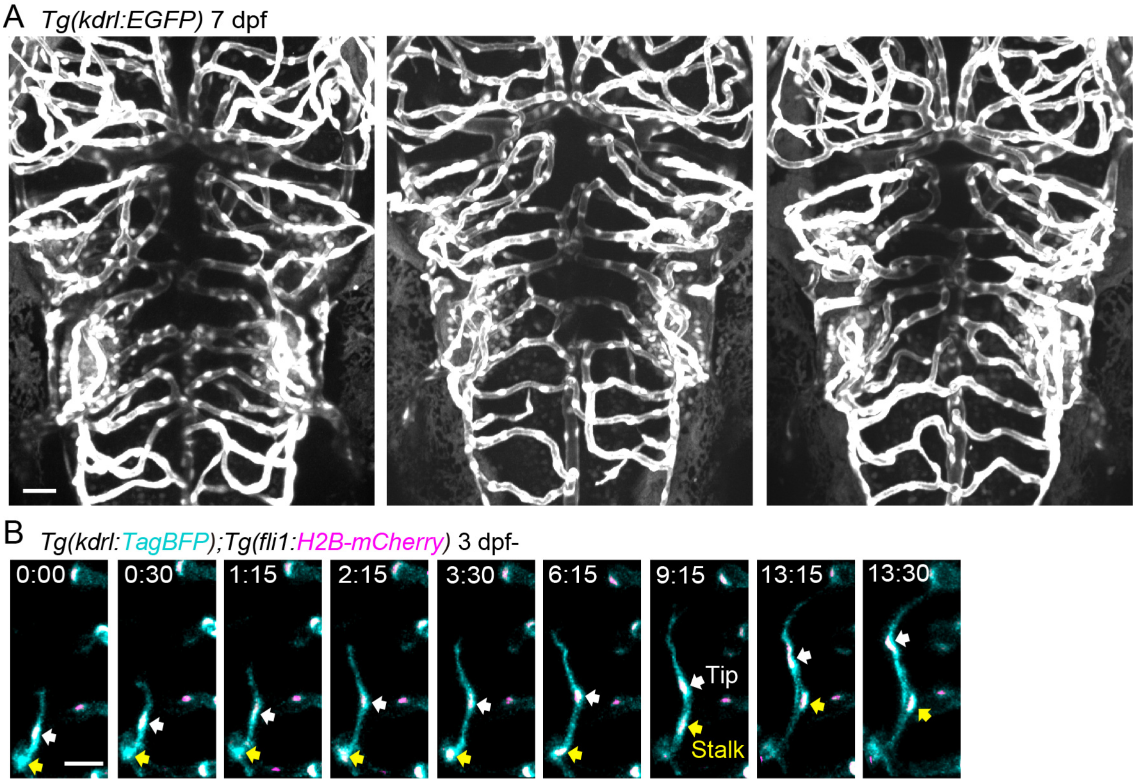


**Figure S1 Formation of capillary networks in the zebrafish brain.**

(A) Representative confocal images of the brain of *Tg(kdrl:EGFP-CAAX)* larvae (7 dpf). *kdrl*:EGFP$^{+}$ endothelial cells (ECs) are shown as white. Dorsal view, anterior to the top. Central artery (CtA) networks show different patterns in different individuals.

(B) Time-sequential images of a *Tg(kdrl:TagBFP)*;*Tg(fli1:H2B-mCherry)* larva (from 3 dpf). Elapsed time (h:min). Tip cell nucleus and stalk cell nucleus are pointed by white arrowheads and yellow arrowheads, respectively. In this sprout, directional migration of the tip cell transiently stopped (2:15 to 9:15) and restarted after the stalk cell (yellow arrowheads) migrated out of the parental vessel.

Scale bar:10 μm.

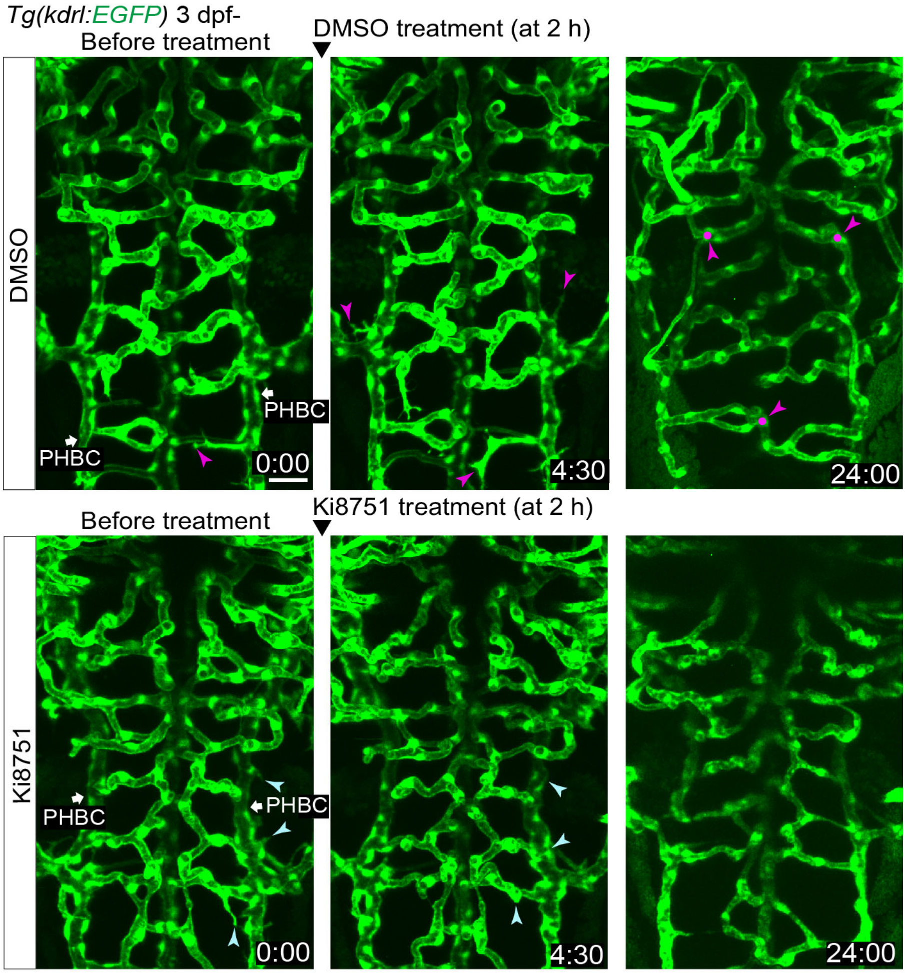


**Figure S2 Inhibition of tip cell sprouting in Vegfr2-inhibited larvae.**

Time-sequential images of *Tg(kdrl:EGFP)* larvae (from 3 dpf) treated with DMSO or 1 μM Ki8751 at 2 h after the start of imaging. Elapsed time (h:min). ECs sprout from the primordial hindbrain channels (PHBC) (magenta arrowheads) and connect with the CtAs (magenta circles) in DMSO-treated larvae, whereas ECs never sprout after Ki8751 treatment. ECs that had already sprouted are retracted soon after Ki8751 treatment (blue arrowheads).

Scale bar:10 μm. PHBC, primordial hindbrain channel.

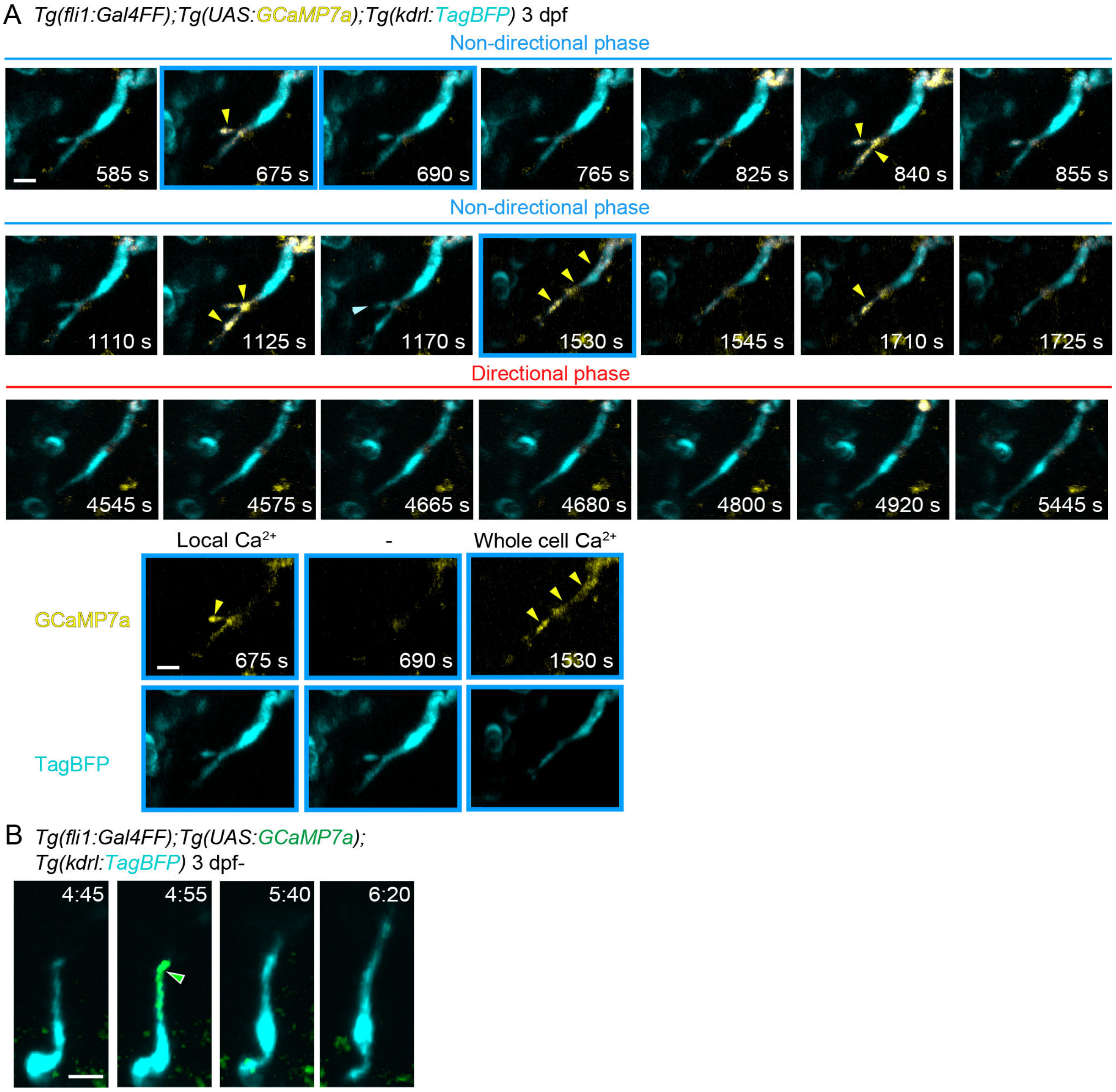


**Figure S3 $Ca^{2+}$ dynamics in tip cells sprouting from the PHBC.**

(A) Time-sequential images of high-speed 3D time-lapse imaging of a *Tg(fli1:Gal4FF);Tg(UAS:GCaMP7a);Tg(kdrl:TagBFP)* larva (3 dpf) taken every 15 sec. In addition to the whole cell $Ca^{2+}$ increase (for example, at 1530 s), local $Ca^{2+}$ oscillations are detected at the leading front of a tip cell in the non-directional phase (yellow arrowheads), but rarely in the directional phase. Elapsed time (sec (s)). The images at 675 s, 690 s, and 1530 s are described in single color as representative of local $Ca^{2+}$ increase, no $Ca^{2+}$ increase, and the whole cell $Ca^{2+}$ increase, respectively (lower panels).

(B) Time-sequential images of a *Tg(fli1:Gal4FF);Tg(UAS:GCaMP7a);Tg(kdrl:TagBFP)* larva (from 3 dpf) after tip cell budding from the PHBC. Local $Ca^{2+}$ increases in an extending protrusion in the non-directional phase (green arrowhead).
Scale bar:10 μm.

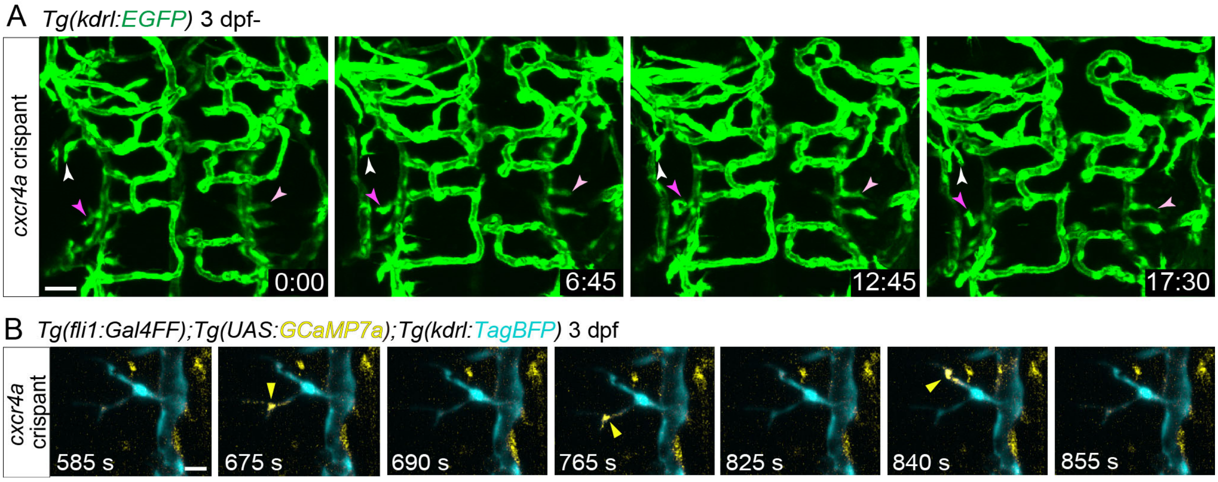


**Figure S4 Phenotypes of *cxcr4a* crispants in cerebral vascular development.**

(A) Time-sequential images of a *Tg(kdrl:EGFP) cxcr4a* crispant larva (F0 injected larva) (3 dpf) after tip cell sprouting from the PHBC. Elapsed time (h:min). Similar to *cxcr4a* mutants, tip cells (arrowheads) do not migrate toward the anastomotic targets in *cxcr4a* crispants.

(B) Time-sequential images of high-speed 3D time-lapse imaging of a *Tg(fli1:Gal4FF);Tg(UAS:GCaMP7a);Tg(kdrl:TagBFP) cxcr4a* crispant larva (3 dpf) taken every 15 sec. Local $Ca^{2+}$ oscillations are detected in protrusions of a tip cell (yellow arrowheads).

Scale bar:10 μm.

### Legends for Supplementary Table and Movies

**Table S1 Sequences of oligonucleotides.**

Sequence list of CRISPR-Cas9 CrRNAs and primers.

**Movie 1 Non-directional and directional migration of a sprouting tip cell from the primordial hindbrain channels (PHBC) toward the central artery (CtA).**

Time-lapse recording in the hindbrain of a *Tg(kdrl:TagBFP);Tg(fli1:H2B-mCherry)* larva (from 3 dpf). Endothelial cells (ECs) and EC nuclei are labeled with TagBFP (cyan) and H2B-mCherry (magenta), respectively. Elapsed time (h:min). Dorsal view, anterior to the top.

**Movie 2 Non-directional migration of a sprouting tip cell from the PHBC in the *cxcr4a* mutant.**

Time-lapse recording in the hindbrain of a *Tg(kdrl:EGFP)* homozygous *cxcr4a* mutant (from 3 dpf). ECs are labeled with EGFP. Elapsed time (h:min). Dorsal view, anterior to the top.

**Movie 3 $Ca^{2+}$ imaging of a sprouting tip cell in the non-directional phase.**

Time-lapse recording in the hindbrain of a *Tg(fli1:Gal4FF);Tg(UAS:GCaMP7a);Tg(kdrl:TagBFP)* larva during the non-directional phase (3 dpf). Elapsed time (sec). Local $Ca^{2+}$ oscillations (yellow) are detected in the leading front of tip cell sprouting from the PHBC. Dorsal view, anterior to the top.

**Movie 4 $Ca^{2+}$ imaging of a sprouting tip cell in the directional phase.**

Time-lapse recording in the hindbrain of a *Tg(fli1:Gal4FF);Tg(UAS:GCaMP7a);Tg(kdrl:TagBFP)* larva during the directional phase (3 dpf). Elapsed time (sec). Local $Ca^{2+}$ oscillations (yellow) are not detected in the leading front of tip cell. Dorsal view, anterior to the top.
